# Leveraging molecular QTL to understand the genetic architecture of diseases and complex traits

**DOI:** 10.1101/203380

**Authors:** Farhad Hormozdiari, Steven Gazal, Bryce van de Geijn, Hilary Finucane, Chelsea J.-T. Ju, Po-Ru Loh, Armin Schoech, Yakir Reshef, Xuanyao Liu, Luke O’Connor, Alexander Gusev, Eleazar Eskin, Alkes L. Price

## Abstract

There is increasing evidence that many GWAS risk loci are molecular QTL for gene expression (eQTL), histone modification (hQTL), splicing (sQTL), and/or DNA methylation (meQTL). Here, we introduce a new set of functional annotations based on causal posterior probabilities (CPP) of fine-mapped molecular cis-QTL, using data from the GTEx and BLUEPRINT consortia. We show that these annotations are very strongly enriched for disease heritability across 41 independent diseases and complex traits (average *N* = 320K): 5.84x for GTEx eQTL, and 5.44x for eQTL, 4.27-4.28x for hQTL (H3K27ac and H3K4me1), 3.61x for sQTL and 2.81x for meQTL in BLUEPRINT (all P ≤ 1.39e-10), far higher than enrichments obtained using standard functional annotations that include all significant molecular cis-QTL (1.17-1.80x). eQTL annotations that were obtained by meta-analyzing all 44 GTEx tissues generally performed best, but tissue-specific blood eQTL annotations produced stronger enrichments for autoimmune diseases and blood cell traits and tissue-specific brain eQTL annotations produced stronger enrichments for brain-related diseases and traits, despite high cis-genetic correlations of eQTL effect sizes across tissues. Notably, eQTL annotations restricted to loss-of-function intolerant genes from ExAC were even more strongly enriched for disease heritability (17.09x; vs. 5.84x for all genes; P = 4.90e-17 for difference). All molecular QTL except sQTL remained significantly enriched for disease heritability in a joint analysis conditioned on each other and on a broad set of functional annotations from previous studies, implying that each of these annotations is uniquely informative for disease and complex trait architectures.

## Introduction

Although Genome-wide association studies (GWAS) have been extremely successful in detecting thousands of risk loci for diseases and traits^1–4^, our understanding of disease architecture is far from complete as most risk loci lie in non-coding regions of genome^5–9,9,10^. Leveraging molecular phenotypes such as gene expression^11–16^ or chromatin marks^17–20^ can aid in understanding disease architecture: in particular, previous studies have shown that cis-eQTL are enriched in GWAS loci as well as genome-wide heritability of several diseases^6,7,21,22^, motivating further work on colocalization^23–28^ and transcriptome-wide association studies^29–31^. Partitioning heritability using raw genotypes/phenotypes^32–36^ or summary association statistics^37–40^ can aid our understanding of disease architectures, but it is currently unclear how to best leverage molecular QTL from rich resources such as GTEx^13,16^ and BLUEPRINT^20^ using these methods.

Here, we introduce a new set of annotations constructed from eQTL, hQTL, sQTL, and meQTL data that are very strongly enriched for disease heritability across 41 independent diseases and complex traits. We construct these annotations by applying a fine-mapping method^41^ (allowing for multiple causal variants at a locus) to compute causal posterior probabilities (CPP) for each variant to be a causal cis-QTL. We show that our annotations are far more enriched for disease heritability than standard annotations that include all significant molecular cis-QTL. We further show that our eQTL annotations produce tissue-specific enrichments (despite high cis-genetic correlations of eQTL effect sizes across tissues^13,42^, and produce much larger enrichments when restricted to loss-of-function intolerant genes from ExAC^43^. Finally, we quantify the extent to which annotations constructed from eQTL, hQTL, sQTL, and meQTL provide complementary information about disease, conditional on each other and on functional annotations from previous studies.

## Results

### Overview of Methods

Our goal is to construct molecular QTL-based annotations that are maximally enriched for disease heritability. For a given molecular QTL data set, we construct a probabilistic (continuous-valued) annotation as follows. First, for each molecular phenotype (e.g. each gene) with at least one significant (FDR < 5%) cis-QTL (e.g. 1Mb from TSS), we compute the causal posterior probability (CPP) of each cis SNP in the fine-mapped 95% credible set, using our CAVIAR fine-mapping method^41^ (see URLs). Then, for each SNP in the genome, we assign an annotation value based on the maximum value of CPP across all molecular phenotypes; SNPs that do not belong to any 95% credible set are assigned an annotation value of 0. We refer to this annotation as MaxCPP. For comparison purposes, we also construct three other molecular QTL-based annotations. First, we construct a binary annotation containing all SNPs that are a significant (FDR < 5%) cis-QTL for at least one molecular phenotype^21,22^; we refer to this annotation as AllcisQTL. Second, we construct a binary annotation containing all SNPs that belong to the 95% credible set (see above) for at least one molecular phenotype; we refer to this annotation as 95%CredibleSet. Third, we construct a binary annotation containing the most significant SNP for each molecular phenotype with at least one significant (FDR < 5%) QTL. We refer to this annotation as TopcisQTL. Further details of each annotation are provided in the Online Methods section.

We apply a previously developed method, stratified LD score regression (S-LDSC)^37,38^, to partition disease heritability using functional annotations. We use two metrics to quantify an annotation’s contribution to disease heritability: enrichment and standardized effect size (τ∗). Enrichment is defined as the proportion of heritability explained by SNPs in an annotation divided by the proportion of SNPs in the annotation^37^; here, we generalize this definition to probabilistic annotations such as MaxCPP. Standardized effect size (τ∗) is defined as the proportionate change in per-SNP heritability associated to a one standard deviation increase in the value of the annotation^38^. Unlike enrichment, τ∗ is conditional on other annotations included in the model. Further details of each metric are provided in the Online Methods section.

We constructed MaxCPP and other annotations using eQTL data from the GTEx Consortium^13,16^ and eQTL, hQTL, sQTL and meQTL data from the BLUEPRINT Consortium^20^ (Table 1; see URLs). We included a broad set of 75 functional annotations from the baselineLD model (Table S1) in most analyses. We have made our annotations freely publicly available (see URLs).

**Table 1.** List of molecular QTL data sets analyzed. GTEx includes eQTL for a wide range of tissues. BLUEPRINT includes eQTL, two hQTL, sQTL and meQTL for 3 blood cell types. Sample sizes for each tissue are provided in Table S5 (GTEx) and Table S25 (BLUEPRINT).

| Dataset | QTL Type | Number of Tissues | $N$ (per tissue) | $N$ (total) |
| --- | --- | --- | --- | --- |
| GTE<sub>x</sub> | eQTL | 44 | 70-361 | 7014 |
| BLUEPRINT | eQTL | 3 | 169-194 | 555 |
| BLUEPRINT | hQTL (H3K27ac) | 3 | 143-174 | 479 |
| BLUEPRINT | hQTL (H3K4me1) | 3 | 104-173 | 449 |
| BLUEPRINT | sQTL | 3 | 169-194 | 555 |
| BLUEPRINT | meQTL | 3 | 132-197 | 525 |

### Simulations

We performed simulations to assess whether S-LDSC produces unbiased estimates of an annotation’s contribution to disease heritability for the AllcisQTL, 95%CredibleSet, TopcisQTL and MaxCPP annotations. The S-LDSC model assumes that causal disease effect sizes are i.i.d. conditional on the value of the annotation, but this assumption may be violated for molecular QTL-based annotations. For example, let x be a SNP in the TopcisQTL annotation, let y be a SNP not in the TopcisQTL annotation that is in LD with x, and let z be a random SNP not in the TopcisQTL annotation. Then, in expectation, y has a larger causal disease effect size than z (violating S-LDSC model assumptions), because it is possible that y is a causal molecular QTL (tagged by x, which may be more significant due to statistical chance), and that y is also causal for disease. The fact that SNPs in LD with the TopcisQTL annotation have larger causal disease effect sizes may cause TopcisQTL enrichment to be overestimated by S-LDSC, which attributes higher χ^2^ statistics for such SNPs entirely to tagging of causal TopcisQTL enrichment. On the other hand, enrichments of more inclusive annotations may be underestimated by S-LDSC, because SNPs in the annotation with high LD to other SNPs in the annotation are expected to have lower causal disease effect sizes than random SNPs in the annotation (again violating S-LDSC model assumptions).

We performed simulations using real genotypes from the UK Biobank, restricting to 749,024 SNPs on chromosome 1 (see Online Methods). We simulated gene expression phenotypes for 500 individuals assuming that each gene has exactly one causal eQTL (heritability=10%), and simulated complex trait phenotypes for an independent set of 40,000 individuals assuming that the set of causal variants is exactly the set of causal eQTLs, with independent effect sizes (heritability=50%). We constructed the AllcisQTL, 95%CredibleSet, TopcisQTL and MaxCPP annotations as described above, and estimated each annotation’s contribution to complex trait heritability using S-LDSC. We performed 400 independent simulations, and averaged results across simulations.

Enrichment estimates and true enrichments for each annotation are displayed in Figure 1 and Table S2. We determined that S-LDSC estimates are severely upward biased for the TopcisQTL annotation, consistent with the above intuition. On the other hand, S-LDSC estimates were conservative for the AllcisQTL, 95%CredibleSet and MaxCPP annotations; thus, we restrict our analyses to these three annotations in the remainder of the manuscript. Of these three annotations, MaxCPP had the highest enrichment (Figure 1 and Table S2) and highest τ∗ (Table S3). We obtained similar results at other values of gene expression heritability and complex trait heritability (Table S4).

**Figure 1.**
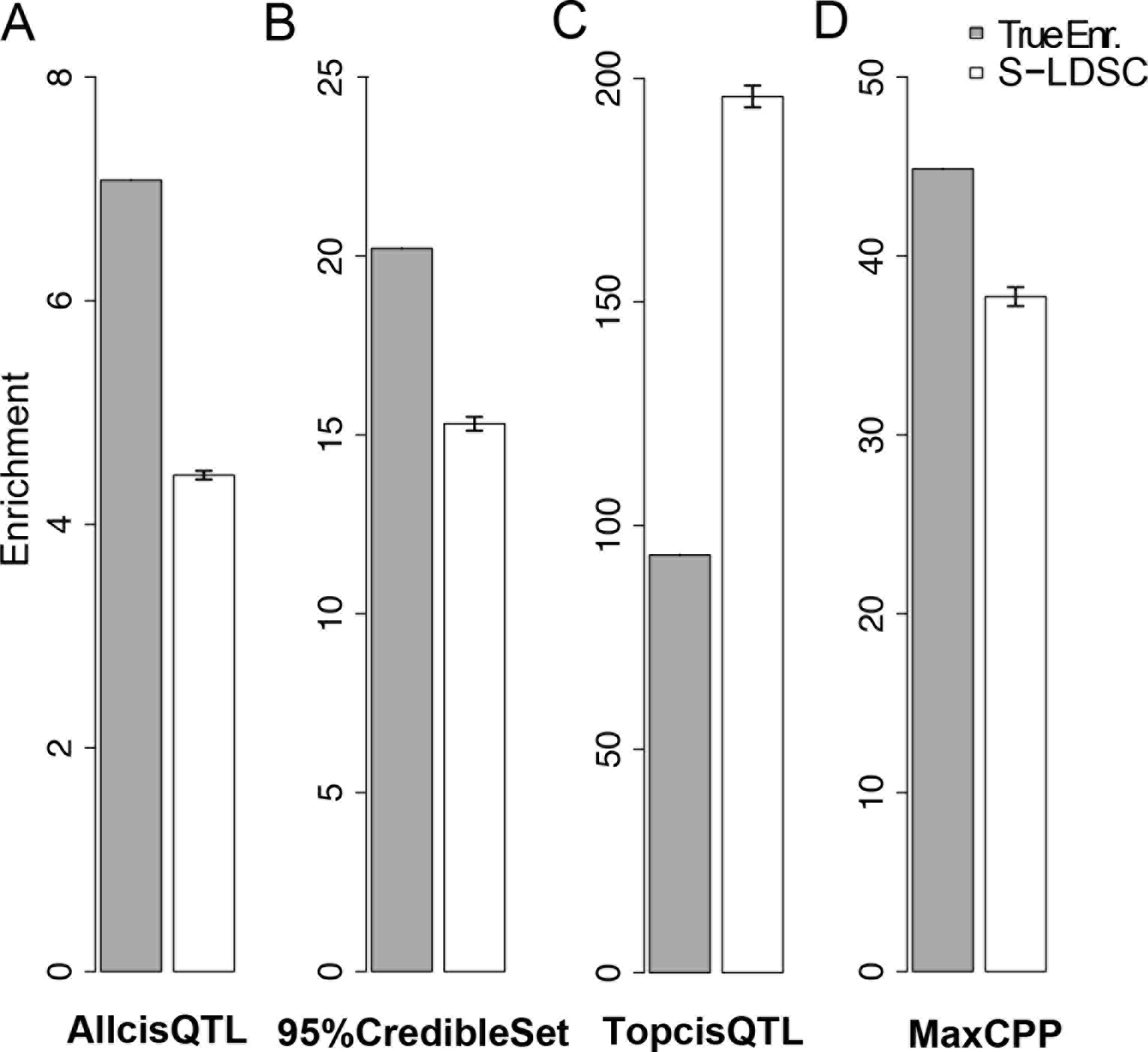
S-LDSC estimates for TopcisQTL are upward biased in simulations. Panels (A), (B), (C), and (D) illustrate the true enrichment and S-LDSC enrichment estimates for AllcisQTL, 95%CredibleSet, TopcisQTL, and MaxCPP annotations, respectively. Error bars represent 95% confidence intervals. Numerical results are reported in Table S2.

### Fine-mapped eQTL are strongly enriched across 41 independent diseases and complex traits

We used the GTEx eQTL data set (Table 1) to construct AllcisQTL, 95%CredibleSet and MaxCPP annotations. We constructed annotations using each of the 44 tissues (Table S5). We applied S-LDSC to assess each annotation’s contribution to disease heritability for each of 41 independent diseases and complex trait data sets (average *N* =320K); for six traits we analyzed two different data sets, leading to a total of 47 data sets analyzed (see Table S6). We meta-analyzed results across the 47 data sets, which were chosen to be independent (no highly genetically correlated traits in overlapping samples; see Online Methods) We computed enrichment and τ∗ for each annotation, in an analysis that included 75 functional annotations (Table S1) from the baselineLD model^38^; we note that enrichment has a simpler interpretation, whereas τ∗ explicitly conditions on annotations from the baselineLD model and is directly comparable across annotations that span different proportions of SNPs. Results for whole blood, a widely studied tissue, are reported in Figure 2 and Table S7. We determined that the MaxCPP annotation had far higher values of enrichment (5.41x, s.e. = 0.82; P = 7.68e-08) and τ∗ (τ∗ = 0.23, s.e. = 0.05; P = 2.00e-05) than AllcisQTL (enrichment = 1.51x and τ∗ = 0.01) or 95%CredibleSet (enrichment = 2.18x and τ∗ = 0.07). The MaxCPP annotation also had higher values of enrichment and τ∗ in analyses that did not condition on the baselineLD model (Figure S1 and Table S8).

**Figure 2.**
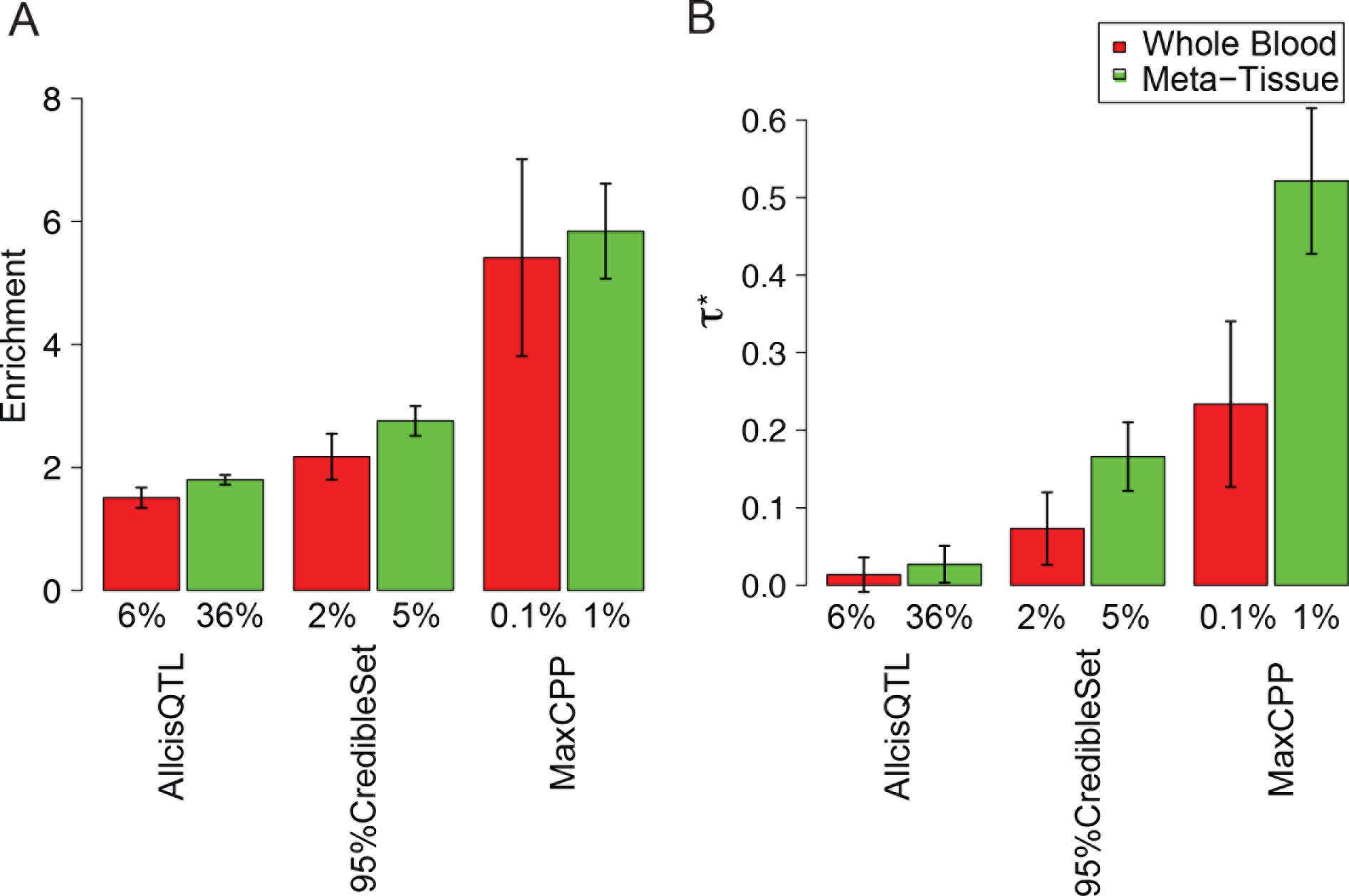
Fine-mapped eQTL are strongly enriched for disease/trait heritability. (A) Meta-analysis results across 41 traits of enrichment for whole blood and Meta-Tissue from the GTEx data set conditioning on the baselineLD model. (B) Meta-analysis results across 41 traits of τ∗ for whole blood and Meta-Tissue conditioning on the baselineLD model. In each panel, we report results for AllcisQTL, 95%CredibleSet, and MaxCPP. Error bars represent 95% confidence intervals. The % value under each bar indicates the proportion of SNPs in each annotation; for probabilistic annotations (MaxCPP), this is defined as the average value of the annotation. Numerical results are reported in Table S7.

We investigated whether the τ∗ for MaxCPP in each respective tissue varied with sample size, which determines the statistical power to detect eQTL. We observed a correlation (*R*^2^ = 0.69, P = 1.36e-12) between sample size and τ∗ (Figure 3 and Table S9). This implies that annotations constructed from tissues with larger sample sizes are more informative for disease/trait architectures.

**Figure 3.**
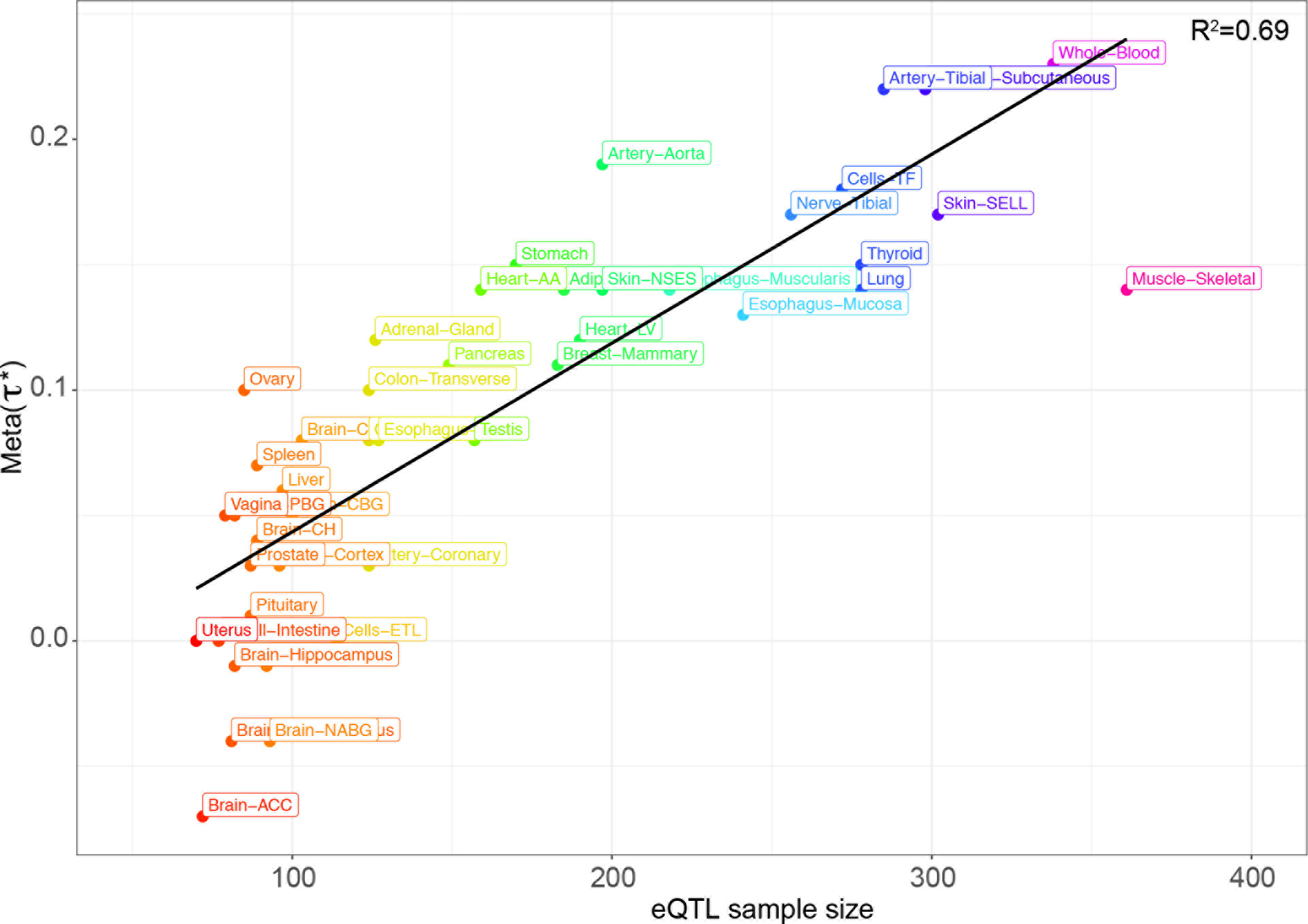
Relationship between eQTL sample size and the annotation effect size (*τ*∗). For each tissue, we plot the τ∗ of the MaxCPP annotation, meta-analyzed across 41 traits, against the eQTL sample size. Numerical results are reported in Table S9. For visualization purposes, we use the following abbreviations: Adipose Visceral Omentum (Adipose-Visceral), Brain Anterior cingulate cortex BA24 (Brain-ACC), Brain Caudate basal ganglia (Brain-CBG), Brain Cerebellar Hemisphere(Brain-CH), Brain Cerebellar Hemisphere (Brain-CH), Brain Frontal Cortex BA9 (Brain-FC), Brain Nucleus accumbens basal ganglia (Brain-NABG), and Brain Putamen basal ganglia (Brain-PBG), Cells EBV transformed lymphocytes (Cells-CETL), Cells Transformed fibroblasts (Cells-TF), Esophagus Gastroesophageal Junction (Esophagus-GJ), Heart Atrial Appendage (Heart-AA), Heart Left Ventricle (Heart-LV), Skin Not Sun Exposed Supra-pubic (Skin-NSES), Skin Sun Exposed Lower leg (Skin-SELL), and Small Intestine Terminal Ileum (Small-Intestine).

To maximize sample size, we performed a fixed-effect meta-analysis of eQTL effect sizes across the 44 tissues (Meta-Tissue); we note that despite pervasive sample overlap, noise is largely uncorrelated across tissues (e.g. average r=0.11 between blood and brain-hippocampus expression despite high cis-genetic correlation of 0.675^42^). We determined that Meta-Tissue annotations had slightly larger enrichments and much larger τ∗ (due to larger annotation size) than annotations constructed from individual tissues (Figure 2, Table S7 and Table S10). As with individual tissues, the MaxCPP annotation had far higher values of enrichment (5.84x, s.e. = 0.40; P = 1.19e-31) and τ∗ (τ∗ = 0.52, s.e. = 0.05; P = 2.73e-27) than AllCisQTL(enrichment = 1.80x and τ∗ = 0.03) or 95%CredibleSet (enrichment = 2.75x and τ∗ = 0.17). A histogram of MaxCPP annotation values for GTEx Meta-Tissue is provided in Figure S2, and correlations with baselineLD model annotations and their LD scores are provided in Figures S3 and S4.

### Tissue-specific fine-mapped eQTL enrichments for blood and brain related traits

Although the Meta-Tissue MaxCPP annotation outperformed each of the 44 tissue-specific MaxCPP annotations in the meta-analysis across 41 traits (Figure 2 and Figure 3), this was not the case for every trait. We examined six autoimmune diseases, five blood cell traits, and eight brain-related diseases and traits in detail (see Online Methods). We first analyzed the six autoimmune diseases, analyzing MaxCPP annotations for blood and Meta-Tissue separately (conditional on the baselineLD model) and meta-analyzing results across the six diseases. We obtained higher estimates of τ∗ (and higher or comparable estimates of enrichment) for blood than for Meta-Tissue or any other individual tissue (Table S11). We then analyzed MaxCPP annotations for blood and Meta-Tissue jointly (conditional on the baselineLD model). We obtained a significantly positive τ∗ estimate for blood (τ∗ = 0.91, s.e. = 0.34; P = 9.15e-03) (Figure 4 and Table S12), implying that fine-mapped blood eQTL provides additional information about these six diseases conditional on fine-mapped Meta-Tissue eQTL. We repeated these analyses for the five blood cell traits. When analyzing MaxCPP annotations for blood and Meta-Tissue separately, we obtained higher estimates of τ∗ (and higher or comparable estimates of enrichment) for blood than for Meta-Tissue or any other individual tissue (Table S13). When analyzing MaxCPP annotations for blood and Meta-Tissue jointly, we obtained a significantly positive τ∗ estimate for blood (τ∗ = 1.17, s.e. = 0.24; P = 1.77e-06) (Figure 4 and Table S12), implying that fine-mapped blood eQTL provides additional information about these five traits conditional on fine-mapped Meta-Tissue eQTL.

**Figure 4.**
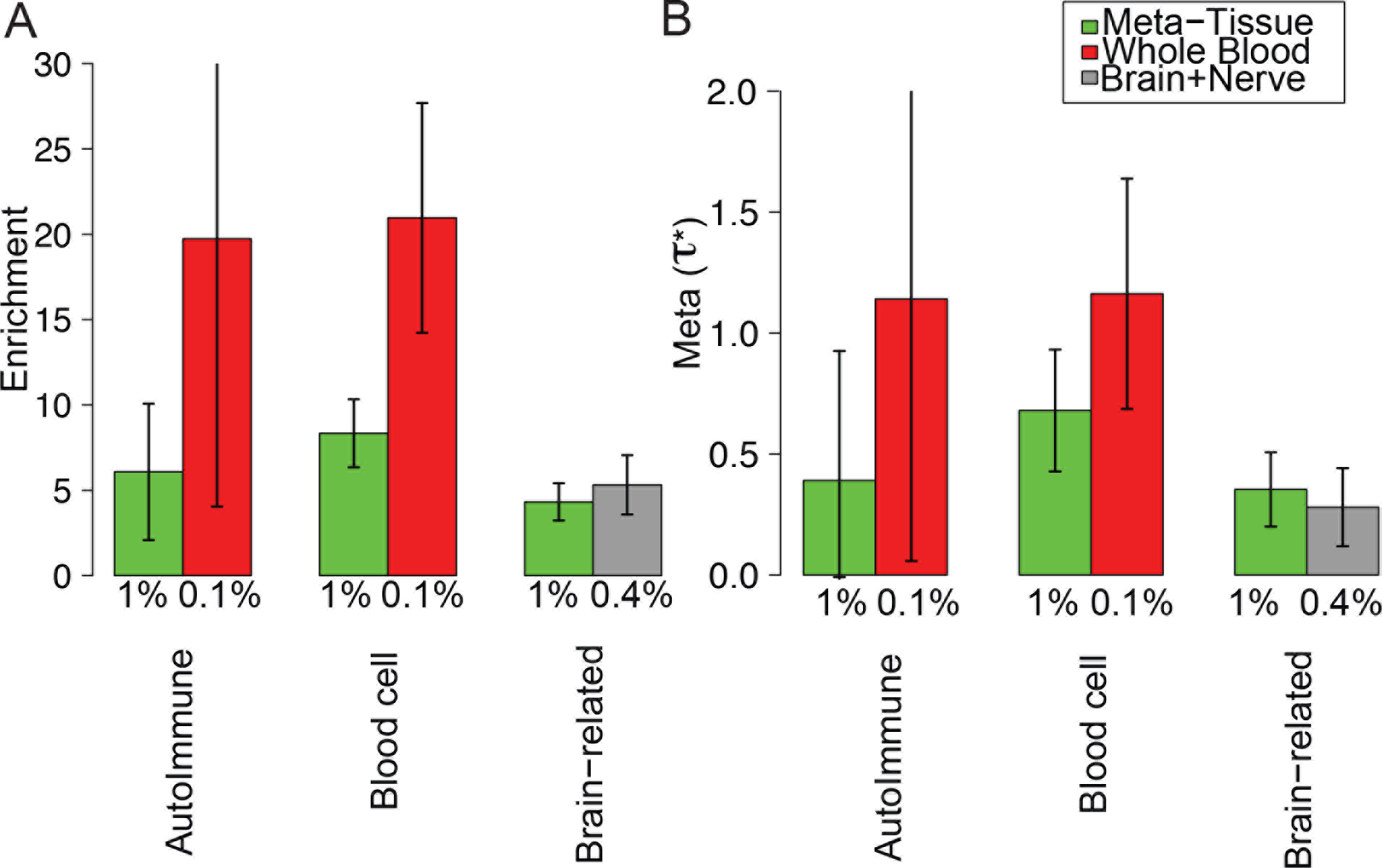
Tissue-specific fine-mapped eQTL enrichments for blood and brain related traits. Meta-analysis results of (A) enrichment and (B) *τ*∗ of Meta-Tissue and tissue-specific MaxCPP annotations, conditional on each other and the baselineLD model, across six independent autoimmune diseases, five blood cell traits, and eight brain-related traits, respectively. Error bars represent 95% confidence intervals. The % value under each bar indicates the proportion of SNPs in each annotation, defined as the average value of the annotation. Numerical results are reported in Table S12.

We then analyzed the eight brain-related diseases and traits. We performed a fixed-effect meta-analysis of eQTL effect sizes across the 10 brain tissues and one nerve tissue (Brain+Nerve). When analyzing MaxCPP annotations for Brain+Nerve and Meta-Tissue separately, we obtained higher or comparable estimates of enrichment and *τ*∗ for Brain+Nerve than for Meta-Tissue or any individual tissue (Table S14). When analyzing MaxCPP annotations for Brain+Nerve and Meta-Tissue jointly, we obtained a significantly positive *τ*∗ estimate for Brain+Nerve (*τ*∗ = 0.28, s.e. = 0.07; P = 9.81e-05) (Figure 4 and Table S12), implying that fine-mapped Brain+Nerve eQTL provides additional information about these eight traits conditional on fine-mapped Meta-Tissue eQTL. We repeated these analyses for each trait and each tissue separately, but determined that that only three blood cell traits (white blood count, red blood cell distribution width, and eosinophil count traits), in conjunction with MaxCPP for blood, attained a significantly positive (FDR<5%) tissue-specific *τ*∗ (Tables S15 and S16). Overall, these results demonstrate that tissue-specific eQTL effects on steady-state expression can be significant for diseases and complex traits, despite the well-documented high cis-genetic correlations of eQTL effect sizes across tissues^13,42^.

### Heritability enrichment of fine-mapped eQTL is concentrated in disease-relevant gene sets

Recent studies have identified gene sets that are depleted for coding variants and enriched for *de novo* coding mutations impacting disease^43–45^. To investigate the importance of non-coding common variants in these gene sets, for each gene set S we used GTEx Meta-Tissue to construct a new annotation MaxCPP(*S*), defined as the maximum CPP restricted to genes in S. For comparison purposes, we also constructed an annotation allSNP(*S*), defined as the set of all SNPs within 100Kb of genes in *S*.

We first analyzed the ExAC gene set, consisting of 3,230 genes that are strongly depleted for protein-truncating variants^43^. We determined that MaxCPP(ExAC) was very strongly enriched in an analysis conditional on the baselineLD model, meta-analyzed across 41 independent traits (see Figure 5A and Table S17). In particular, MaxCPP(ExAC) was much more strongly enriched (17.06x, s.e. = 1.28; P = 1.20e-35) than MaxCPP(All Genes) (5.84x) (P = 4.90e-17 for differ-ence). This implies that eQTL for these 3,230 genes have a disproportionately strong impact on disease heritability, consistent with the fact that these genes are depleted for eQTL^43^. We then analyzed MaxCPP(ExAC) and MaxCPP(All Genes) annotations jointly (conditional on the baselineLD model). We obtained a significantly positive *τ*∗ for MaxCPP(ExAC) (*τ*∗ = 0.42, s.e. = 0.04; P = 1.40e-23; Figure 5B and Table S18), implying that MaxCPP(ExAC) provides additional information about disease heritability conditional on MaxCPP (All Genes). We observed that the effect size (*τ*∗) for MaxCPP(ExAC) conditional on MaxCPP (All Genes) and baselineLD is five times larger, and more statistically significant, than the *τ*∗ of allSNP (ExAC) conditional on the baselineLD model (Table S24). Thus, MaxCPP can increase power to identify enriched gene sets.

**Figure 5.**
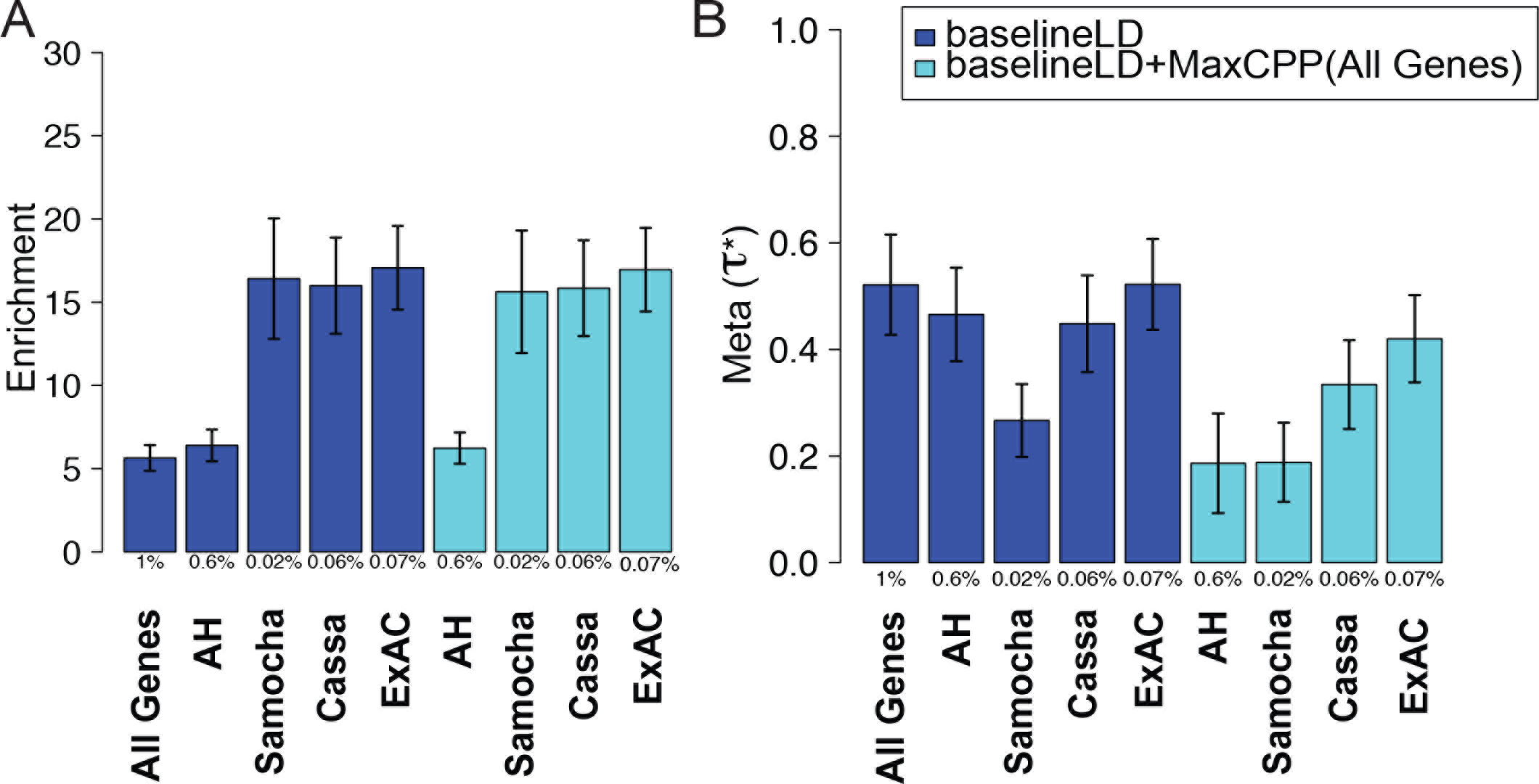
Heritability enrichment of fine-mapped eQTL is concentrated in disease-relevant gene sets. Meta-analysis results of (A) enrichment and (B) *τ*∗ of MaxCPP(S) for various gene sets S. We report results conditional on the baselineLD model (dark blue) and results conditional on both the baselineLD model and MaxCPP(All Genes) (light blue), meta-analyzed across 41 traits. As expected, *τ*∗ estimates are reduced by conditioning on MaxCPP(All Genes), but enrichment estimates are not affected. Error bars represent 95% confidence intervals. The % value under each bar indicates the proportion of SNPs in each annotation, defined as the average value of the annotation. Numerical results are reported in Table S17.

We analyzed four additional gene sets S: a set of 1,003 genes that are strongly depleted for missense mutations^44^ (Samocha); a set of 2,984 genes with strong selection against protein-truncating variants^45^ (Cassa); a set of 1,878 genes predicted to be essential based on CRISPR experiments in a human cancer cell line^46^ (Wang); and a set of 11,983 genes with evidence of allelic heterogeneity in analyses of GTEx gene expression data using our previously developed methods (AH)^47^. For each of these gene sets, MaxCPP(S) was strongly enriched in analyses conditional on the baselineLD model, meta-analyzed across 41 independent traits (Figure 5A and Table S17). In addition, for each gene set except the Wang gene set, we obtained a significantly positive *τ*∗ for maxCPP(S) (after correcting for five gene sets tested) when analyzing MaxCPP(S) and MaxCPP(All Genes) jointly (conditional on the baselineLD model) (Figure 5B, Table S18 and Tables S19-S23). As with the ExAC gene set, the *τ*∗ for MaxCPP(S) conditional on MaxCPP(All Genes) and the baselineLD model were substantially larger than the *τ*∗ of allSNPs(ExAC) conditional on the baselineLD model and were often more statistically significant (Table S24), indicating that MaxCPP can increase power to identify enriched gene sets in which regulatory variants play an important role.

### Fine-mapped QTL for a broad set of molecular traits are strongly enriched across 41 independent diseases and complex traits

We analyzed five molecular QTL from the BLUEPRINT data set (Table 1), including eQTL, hQTL (H3K27ac and H3K4me1), sQTL and meQTL. In each case, we constructed AllcisQTL, 95%CredibleSet and MaxCPP annotations using each of the three immune cell types (CD14+ monocytes, CD16+ neutrophils, and naive CD4+ T cells; Table S25) as well as a fixed-effect meta-analysis of molecular QTL effect sizes across the 3 cell types (Meta-Tissue). We again used S-LDSC to assess each annotation’s contribution to disease heritability, meta-analyzed across 41 independent diseases and complex traits (47 data sets). We determined that for each QTL data set the MaxCPP annotation (enrichment = 2.06-6.02x and *τ*∗ = 0.02-0.45) outperformed the AllcisQTL (enrichment = 1.17-1.56x and *τ*∗ = 0.01-0.09) and 95%CredibleSet (enrichment = 1.44-2.28x and *τ*∗ = −0.01-0.16) annotations (Table S26). A histogram of MaxCPP annotation values for each QTL data set is provided in Figure S5. MaxCPP for each molecular QTL was significantly enriched in an analysis conditional on the baselineLD model, meta-analyzed across the 41 traits: eQTL (5.44x, s.e. = 0.55; P = 3.26e-16), H3K27ac (4.28x, s.e. = 0.37; P = 2.59e-19), H3K4me1 (4.27x, s.e. = 0.36; P = 1.29e-20), sQTL (3.61x, s.e. = 0.40; P =1.39e-10), and meQTL (2.81x, s.e. = 0.19; P = 8.36e-22) (Figure 6A and Table S27). Meta-Tissue generally attained higher enrichments and *τ*∗ than MaxCPP computed using each of the three immune cell types individually (Table S28), similar to our GTEx results (Tables S7 and S9). MaxCPP computed using Meta-Tissue also generally outperformed each of the three cell types in a meta-analysis across the six autoimmune diseases (Table S29) and a meta-analysis across the five blood cell traits (Table S30), in contrast to the stronger enrichments for tissue-specific GTEx blood eQTL annotations for blood cell traits (Tables S11 and S13).

**Figure 6.**
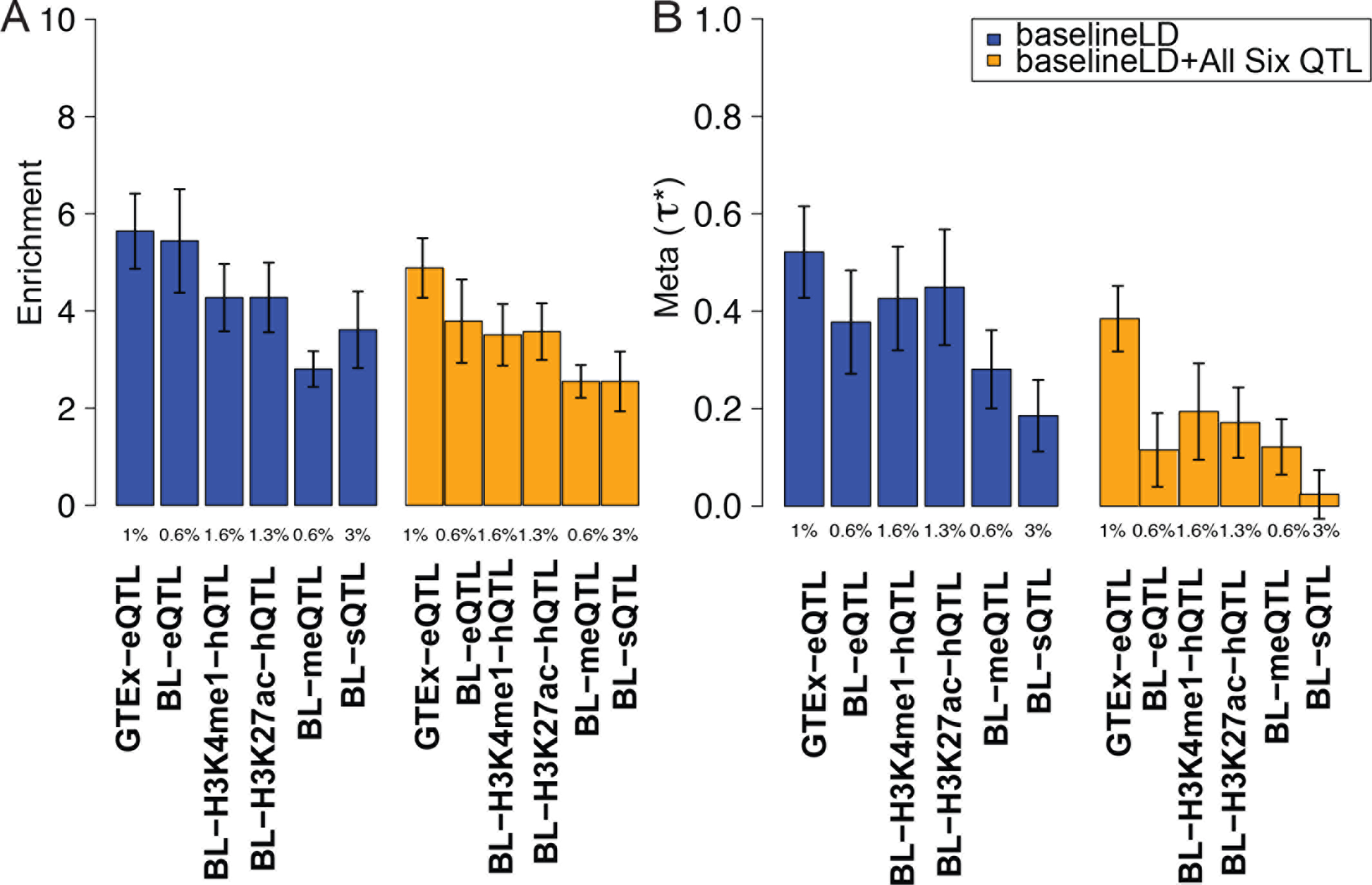
Fine-mapped eQTL, hQTL, sQTL, and meQTL annotations are enriched for dis-ease/trait heritability. Meta-analysis results of (A) enrichment and (B) *τ*∗ of MaxCPP for various molecular QTL from GTEx and BLUEPRINT (BL). We report results conditional on the baselineLD model (dark blue) and results conditional on both the baselineLD model and MaxCPP for all six molecular QTL (orange), meta-analyzed across 41 traits. As expected, *τ*∗ estimates are reduced by conditioning on Max-CPP for all molecular QTL, but enrichment estimates are not affected. Error bars represent 95% confidence intervals. The % value under each bar indicates the proportion of SNPs in each annotation, defined as the average value of the annotation. Numerical results are reported in Table S27 and Table S31.

Finally, we jointly analyzed MaxCPP annotations for GTEx eQTL and each of the five BLUEPRINT molecular QTL (conditional on the baselineLD model). The purpose of this analysis was to determine whether each of these molecular QTL provides independent information about disease and complex trait architectures. We determined that *τ*∗ remained statistically significant for all molecular QTL except sQTL (Figure 6B and Table S31); a joint analysis of just the five BLUEPRINT molecular QTL (conditional on the baselineLD model) produced similar findings (Table S32). LD scores of the sQTL annotation had the highest correlation with LD scores of the GTEx eQTL and BLUEPRINT eQTL annotations (*R* = 0.56 −0.57; see Figure S4), implying that much of the informativeness of sQTL in this analysis is captured by eQTL. However, eQTL, hQTL (H3K27ac and H3K4me1) and meQTL are each uniquely informative for disease and complex trait architectures.

## Discussion

We have shown that annotations constructed using fine-mapped posterior probabilities for several different molecular QTL are very strongly enriched for disease heritability across 41 diseases and complex traits. These results improve upon two previous studies that made key contributions in showing that annotations constructed using all significant cis-eQTL were significantly enriched for heritability of obsessive-compulsive disorder and Tourette Syndrome^21^ as well as type 2 diabetes^22^. Our findings provide additional motivation for colocalization studies^23–27^ to identify shared causal variants for eQTL and disease and transcriptome-wide association studies (TWAS)^29–31^ to identify genes whose predicted expression is associated to disease. Our fine-mapped eQTL annotations were able to detect tissue-specific enrichments for blood and brain related traits, despite high cis-genetic correlations^13,42^ of eQTL effect sizes across tissues and despite the fact that TWAS have generally concluded that their results “did not suggest tissue-specific enrichment”^31^.

We note that a previous study showed that cis-eQTL often lie close to the transcription start site (TSS) or transcription end site (TES)^48^; we did not observe additional disease signal for cis-eQTL variants close to the TSS or TES in the GTEx or BLUEPRINT data sets (see Tables S33 and S34). Notably, our eQTL annotations produced particularly large enrichments when restricted to disease-relevant gene sets such as loss-of-function intolerant genes from ExAC, highlighting the potential to increase signal in analyses of gene sets harboring regulatory signals by prioritizing fine-mapped cis-eQTL.

We also detected strong enrichments using annotations based on other molecular QTL, with eQTL, hQTL and meQTL all providing complementary information about disease, conditional on each other and on functional annotations from previous studies. These results motivate applying colocalization and TWAS methods to other molecular QTL; it may also be possible to prioritize other molecular QTL in gene set analyses by connecting regulatory regions to genes^49–53^. We note that although annotations constructed from sQTL were not conditionally significant in our analysis, previous work has shown that sQTL can contain information that is independent from eQTL^54^, motivating further investigation in larger sQTL data sets. Finally, although we have used the CAVIAR method^41^ for fine-mapping molecular QTL, our results motivate a comparison of disease/trait enrichment using other QTL fine-mapping methods^55–59^.

We note several limitations of our work. First, we restrict our analyses to common variants, as S-LDSC is not currently applicable to rare variants^37^. Recent work has shown that rare variants can have substantial effects on gene expression^60^, motivating ongoing work to extend S-LDSC to rare variants. Second, the CAVIAR fine-mapping method allows up to six causal variants per locus; this may limit power at loci that harbor more than six causal variants, although this would not lead to spurious signals. Third, we show that S-LDSC is generally unable to produce unbiased enrichment estimates for molecular QTL based annotations, which violate its model assumptions; S-LDSC produces slightly conservative enrichment estimates for the MaxCPP annotation that we focus on here, although we caution that the TopcisQTL annotation produces large upward biases and should be avoided (Figure 1). Fourth, our results are a function of the molecular QTL sample size (Figure 3) and set of tissues; analyses of larger sample sizes and/or different tissues or contexts may produce larger enrichments in the future. Fifth, although we analyzed very large GWAS samples sizes (average N=320K), power increases only modestly with GWAS sample size at very large sample sizes, as the finite size of the genome is a stricter constraint^61^. Sixth, we performed a fixed-effect meta-analysis of molecular QTL effect sizes across tissues (Meta-Tissue), but performing fixed-effect meta-analysis with overlapping samples can limit power^62–64^. We believe that this is a minor issue in the current context, as noise is largely uncorrelated across tissues, but further investigation using random-effect meta-analysis is warranted. Despite these limitations, our results indicate that fine-mapped QTL annotations are strongly enriched for disease heritability and can help elucidate the genetic architecture of diseases and complex traits.

## Acknowledgements

We are grateful to Soumya Raychaudhuri, Noah Zaitlen, Bogdan Pasaniuc, and Fereydoun Hormozdiari for helpful discussions. This research was funded by NIH grants U01 HG009379, R01 MH101244, R01 MH109978 and R01 MH107649. This research was conducted using the UK Biobank Resource under Application 16549.

## Online Methods

### Molecular QTL-based annotations

We construct four annotations for any given QTL data set using the observed marginal association statistics. The four annotations are MaxCPP, AllcisQTL, 95%CredibleSet, and TopcisQTL. Each annotation is a vector that assigns a value to each SNP. Let a indicate our annotation for one QTL data set where *a*_*j*_ indicates the value assigned to SNP *j*. For binary annotations (AllcisQTL, 95%CredibleSet, and TopcisQTL) *a*_*j*_ ∈ {0, 1}, and *a*_*j*_ = 0 indicates that SNP *j* is not included in the annotation, while *a*_*j*_ = 1 indicates that SNP *j* is included in the annotation. For continuous probabilistic annotations (MaxCPP), 0 ≤ *a*_*j*_ ≤ 1.

#### MaxCPP annotation

Let ***S* = (*s*_1_, *s*_2_, … *s*_*g*_)** indicate an (m ×g) matrix of the observed marginal association statistics obtained for each QTL data set, where m is the number of SNPs and g is the number of eGenes (e.g., genes that have at least one significant cis-eQTL). Let *s*_*i*_ be the vector of marginal association statistics of gene *i* for all the cis-variants. Utilizing *s*_*i*_ and the LD structure, we can compute the causal posterior probability (CPP) for each variant. CPP is the probability that a variant is causal. Let *α*_*ji*_ be the posterior probability that the SNP *j* is causal for the gene *i*. We obtain the CPP values from CAVIAR^41^. In addition to the CPP values, CAVIAR provides a 95%credible set that contains all of the causal variants with probability at least 95%. Let *θ*_*ji*_ indicate whether SNP j is in the 95%credible set for the gene *i* (i.e., *θ*_*ji*_=1 indicates that the SNP *j* is in the gene *i* 95%credible set and *θ*_*ji*_=0 otherwise). We construct the MaxCPP annotation for SNP *j* by computing the maximum value of CPP over all genes where SNP *j* is in the 95%credible set of the gene *i*. More formally, we have: *a*_*j*_ = max_*i*_ *α*_*ji*_ where the maximum is over genes *i* with *θ*_*ij*_ = 1.

#### AllcisQTL annotation

AllcisQTL annotation is a binary annotation, where any variant whose marginal association statistic for at least one gene passes the significance threshold (FDR < 0.05) has annotation value 1, and each other variant has annotation value 0. Let t_ji_ indicate whether the SNP j is statistically significant for the gene *i* (*t*_*ji*_ = 1 when FDR(j) < 0.05 and *t*_*ij*_ = 0 otherwise). More formally, we have: *a*_*j*_ = max_*i*_ *t*_*ji*_.

#### 95%CredibleSet annotation

95%CredibleSet is a binary annotation, any variant that is in a 95%credible set of at least one gene has annotation value 1 and each other variant has annotation value 0. More formally, we have: *a*_*j*_ = max_*i*_ *θ*_*ji*_.

#### TopcisQTL annotation

TopcisQTL is a binary annotation where any variant that is the most significant variant for at least one gene has annotation value 1 and each other variant has annotation value 0. Let γ_*ji*_ indicate whether the SNP *j* is the most significant SNP for the gene *i* (i.e., γ_*ji*_=1 if SNP *j* is the most significant SNP among all cis-variants for the gene i and γ_*ji*_=0 otherwise). More formally, we have: *a*_*j*_ = max_*i*_ γ_*ji*_.

### Enrichment and effect size (*τ*∗) metrics

We use two metrics to measure the importance of an annotation in the context of diseases and complex traits: Enrichment and effect size (*τ*∗) of annotation. We use S-LDSC^37,38^ to compute enrichment and effect size (*τ*∗). Let a_cj_ indicate the annotation value of the SNP *j* for the annotation *c*. S-LDSC^37,38^ assumes that the variance of each SNP is a linear additive contribution to each annotation:

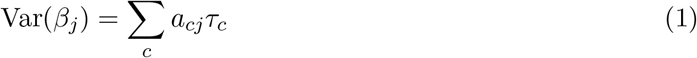

where *τ*_*c*_ is the contribution of the annotation *c* to per-SNP heritability. S-LDSC^37,38^ estimates *τ*_c_ using the following equation:

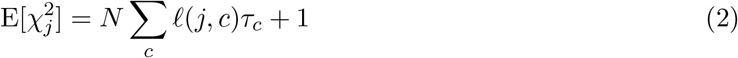

where *N* is the GWAS sample size and *ℓ*(*j*, *c*) is the LD score of the SNP *j* for the annotation *c*. S-LDSC computes the LD scores as follow: 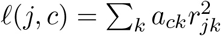 where *r*_*jk*_ is the genetic correlationbetween the SNPs *j* and *k*.

Since *τ*_*c*_ depends on the trait heritability and the size of the annotation, ref.^38^ defined *τ*_*c*_ for an annotation as the per-standardized annotation effect size:

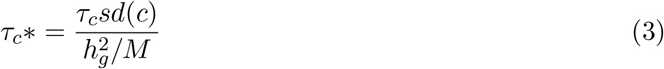

where *sd*(*c*) is the standard deviation of the annotation *c*, 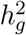 is the SNP-heritability, and *M* is the number of variants used to compute 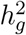. In our experiments, *M* is equal to 5,961,159 (see below).

The enrichment of an annotation is defined as the fraction of heritability captured by the annotation divide by the fraction of SNPs in that annotation. We extend the definition of enrichment to continuous probabilistic annotations with values between 0 and 1:

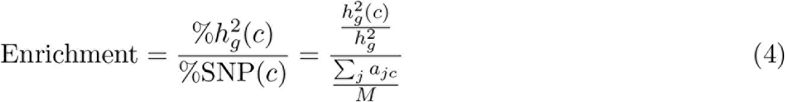

where 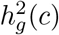 is the heritability captured by the *c*-th annotation. We can compute this quantity as follows:

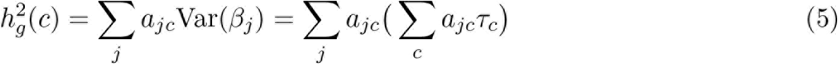

Although enrichment has a simpler interpretation than *τ*_*c*_×, *τ*_*c*_× is computed by conditioning on other annotations in the model. Thus, *τ*_*c*_× captures the signal that is unique to the annotation c and is not captured by other annotations in the model. For example, two annotations that are highly correlated will have similar enrichment, while *τ*∗ for the annotation that better captures true causal variants will be larger and more statistically significant.

### Simulation framework

In our simulation, we simulated both gene expression and trait phenotypes. We utilized UK Biobank genotypes from chromosome 1, which consists of 749,024 variants, for our simulation. We used 40,000 individuals to generate the trait phenotypes and a non-overlapping set of 500 individuals to generate gene expression phenotypes. We implanted one causal variant per each gene for all the 4,493 genes on chromosome 1, and assumed that the set of causal trait variants was exactly the same set of 4,493 variants.

We set the total heritability for each gene and trait phenotype to be 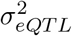 and 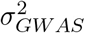, respectively. We sisimulated causal trait e.ect sizes using polygenetic model, 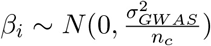, where *β*_2_ is the true effect size of *i*-th causal variant and *n*_*c*_ is the number of causal variants. In the case of gene expression, we simulated 4,493 phenotypes to represent our simulated gene expression data and we set *n*_*c*_ to one for each gene. In the case of trait phenotypes, we set *n*_*c*_ to 4,493 as the number of causal variant is equal to the number of genes in chromosome 1. In our simulated data sets, we set 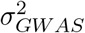 to 0.5 and considered different values of 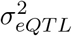 to test our results on different input parameters. We set 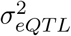 to 0.05, 0.1, and 0.2 resulting in three different simulated data sets. For each simulated data set, we performed 400 simulations.

After simulating the gene expression and trait phenotypes, we obtained marginal association statistics for each variant using linear regression implemented in the PLINK software^65^ (see URLs). In the case of simulated trait phenotypes, we computed the association of each variant with the simulated trait phenotypes. In the case of simulated gene expression phenotypes, we computed the marginal statistics for all variants within 1Mb of the TSS. We generated the four annotations as described above. We used CAVIAR^41^ to generate the 95%Credible set and MaxCPP annotations. We utilized European samples from the 1000 Genomes Project (1000G)^66^ (see URLs) to estimate the LD structure required as input to CAVIAR. We applied CAVIAR under a setting where we allowed up to six causal variants for each gene.

After obtaining the four annotations, we ran S-LDSC^37^ to generate the LD score of each variant in each annotation using the same procedure described in the previous studies^37,38^. Regression SNPs, which are used by S-LDSC to estimate *τ* from marginal association statistics, were obtained from the HapMap Project phase 3^67^. These SNPs are considered as well-imputed SNPs. SNPs with marginal association statistics larger than 80 or larger than 0.001 *N* and SNPs that are in the major histocompatibility complex (MHC) region were excluding from all the analyses. Reference SNPs, which are used to compute LD scores, were defined using the European samples in 1000G^66^. Heritability SNPs, which are used to estimate *sd*(*c*) and 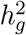, were defined as common variants (MAF ≥ 0.05) in the set of reference SNPs. Using the LD score for each annotation and the marginal statistics obtained from the trait phenotypes, we computed the enrichment and *τ*∗ for each simulation. Then, we compared the S-LDSC estimated enrichment and true enrichment for each annotation. All results are averaged across 400 simulations.

### Set of 41 independent diseases and complex traits

We initially considered 34 GWAS summary association statistic data sets that are publicly available and 55 UK Biobank traits (see URLs) for which which summary association statistics were computed using BOLT-LMM (see URLs; up to N=459K European-ancestry samples). We restricted our analyses to 47 data sets with z-scores of total SNP heritability at least 6 (Table S6). The 47 data sets included 6 traits that were duplicated in two different data sets (genetic correlation of at least 0.9, Table S6). Thus, we analyzed 41 independent diseases and complex traits. We ran S-LDSC using the same procedures described in previous studies^37,38^ (see above). Most analyses included the baselineLD model v1.1, which is identical to the baselineLD model as previously described^38^ except that we fixed an error in the promoter annotation (inherited from previous studies^34,37^); we determined that fixing this error did not affect our results. The z-score of total SNP heritability was computed using S-LDSC with the baselineLD model, and the genetic correlation between pairs of traits was computed using cross-trait LDSC^69^. The meta-analyzed values of enrichment and *τ*∗ across the 47 data sets were computed using a random-effect meta-analysis, as implemented in the rmeta R package.

### Blood and brain related diseases and complex traits

#### Autoimmune diseases

We analyzed six autoimmune diseases: Crohn’s disease^70^, Rheumatoid arthritis (ref.^71^ and UK Biobank), Ulcerative colitis^70^, Lupus^72^, Celiac^73^, and all autoimmune and inflammatory diseases in UK Biobank).

#### Blood cell traits

We analyzed five blood cell traits: white blood cell count, red blood cell count, platelet count, eosinophil count, and red blood cell distribution width. All of these data sets were obtained from UK Biobank.

#### Brain-related diseases and complex traits

We analyzed eight independent brain-related diseases and complex traits: Age at menarche^74^, BMI (ref.^75^ and UK Biobank), Bipolar Disorder^76^, Depressive symptoms^3^, Neuroticism (UK Biobank), Schizophrenia^1^, Smoking Status (ref.^77^ and UK Biobank), and Year of education (ref.^78^ and UK Biobank). These traits are a subset of traits from Table S6 that were reported to be brain-enriched^37,61^.

## URLs

CAVIAR: http://genetics.cs.ucla.edu/caviar/
GTEx (Release v6, dbGaP Accession phs000424.v6.p1): http://www.gtexportal.org.
BLUEPRINT: ftp://ftp.ebi.ac.uk/pub/databases/blueprint/blueprintEpivar/qtlas/
baselineLD annotations: https://data.broadinstitute.org/alkesgroup/LDSCORE/
QTL-based annotations: https://data.broadinstitute.org/alkesgroup/LDSCORE/
1000 Genomes Project Phase 3 data: ftp://ftp.1000genomes.ebi.ac.uk/vol1/ftp/release/20130502
PLINK software: https://www.cog-genomics.org/plink2
BOLT-LMM software: https://data.broadinstitute.org/alkesgroup/BOLT-LMM
BOLT-LMM summary statistics for UK Biobank traits: https://data.broadinstitute.org/alkesgroup/UK
UK Biobank: http://www.ukbiobank.ac.uk/
UK Biobank Genotyping and QC Documentation: http://www.ukbiobank.ac.uk/wp-content/uploads/201404/UKBiobankgenotypingQCdocumentation-web.pdf

